# Sensorimotor functional connectivity in unilateral cerebral palsy: influence of corticospinal tract wiring pattern and clinical correlates

**DOI:** 10.1101/515866

**Authors:** Cristina Simon-Martinez, Ellen Jaspers, Kaat Alaerts, Els Ortibus, Joshua Balsters, Lisa Mailleux, Jeroen Blommaert, Charlotte Sleurs, Katrijn Klingels, Frédéric Amant, Anne Uyttebroeck, Nicole Wenderoth, Hilde Feys

**Affiliations:** KU Leuven Department of Rehabilitation Sciences, Leuven, Belgium; Neural Control of Movement Lab, Department of Health Sciences and Technology, ETH Zurich, Switzerland; KU Leuven Department of Development and Regeneration, Leuven, Belgium; Department of Psychology, Royal Holloway University of London, Egham, United Kingdom; KU Leuven Department of Oncology, Leuven, Belgium; Rehabilitation Research Centre, Faculty of Rehabilitation Sciences, Hasselt University, Diepenbeek, Belgium; Centre for Gynaecologic Oncology, Antoni van Leeuwenhoek, Amsterdam, Netherlands; Centre for Gynaecologic Oncology, Amsterdam University Medical Centres, Amsterdam, Netherlands

**Keywords:** cerebral palsy, functional neuroimaging, paediatrics, sensorimotor cortex, upper extremity

## Abstract

In children with unilateral cerebral palsy (uCP), the corticospinal tract (CST) wiring patterns may differ (contralateral, ipsilateral or bilateral), partially determining motor deficits. However, the impact of such CST wiring on functional connectivity remains unknown. Here, we explored differences in functional connectivity of the resting-state sensorimotor network in 26 uCP with periventricular white matter lesions (mean age (SD): 12.87m (±4.5), CST wiring: 9 contralateral, 9 ipsilateral, 6 bilateral) compared to 60 healthy controls (mean age (SD): 14.54 (±4.8)), and between CST wiring patterns. Functional connectivity from each M1 to three bilateral sensorimotor regions of interest (primary sensory cortex, dorsal and ventral premotor cortex) and the supplementary motor area was compared between groups (healthy controls vs. uCP; and healthy controls vs. each CST wiring group). Results from the seed-to-voxel analyses from bilateral M1 were compared between groups. Additionally, relations with upper limb motor deficits were explored. Aberrant sensorimotor functional connectivity seemed to be CST-dependent rather than specific from all the uCP population: in the dominant hemisphere, the contralateral CST group showed increased connectivity between M1 and premotor cortices, whereas the bilateral CST group showed higher connectivity between M1 and somatosensory association areas. These results suggest that functional connectivity of the sensorimotor network is CST wiring-dependent, although the impact on upper limb function remains unclear.

## 1. Introduction

Upper limb (UL) function is commonly impaired in individuals with unilateral cerebral palsy (uCP), negatively influencing the performance of daily life activities (Klingels et al., 2012). Given the large variability in clinical presentation of UL function, there has been increasing interest in investigating the underlying neural mechanisms, with the aim of developing individually targeted rehabilitation programs.

After brain injury, different neuroplastic mechanisms take place and the resulting functional reorganization may not necessarily correspond with the remaining structural connectivity. Several efforts have been made to investigate the underlying pathophysiology of uCP or upper limb impairments by targeting the structural properties of the brain injury. These studies suggest that both structural brain lesion characteristics and microstructural integrity of the white matter bundles partially explain the variability in UL dysfunction. More specifically, later and larger lesions and lower integrity of cortico-subcortical tracts lead to worse function (Feys et al., 2010; L. Holmström et al., 2011; Mailleux et al., 2017; Tsao, Pannek, Fiori, Boyd, & Rose, 2014). Moreover, the underlying corticospinal tract (CST) wiring has been put forward as an important explanatory factor (Gupta et al., 2017; Linda Holmström et al., 2010; Simon-Martinez, Jaspers, et al., 2018; Staudt, 2010; Zewdie, Damji, Ciechanski, Seeger, & Kirton, 2017), indicating that children with a contralateral CST wiring have a more preserved motor function than those with bilateral or ipsilateral CST wiring. Although the type of CST wiring seems to be a relevant biomarker of motor function, we recently have shown large variability in UL deficits within the bilateral and ipsilateral CST groups (Simon-Martinez, Jaspers, et al., 2018), which might be better explained by how these groups functionally integrate different brain areas of the sensorimotor network to execute arm and hand movements.

The relation between functional connectivity and UL function in uCP has mainly been studied using task-based functional MRI (fMRI) (Gaberova, Pacheva, & Ivanov, 2018). However, considerable inter-study variability regarding task choice and the dependence on the ability of the child to adequately perform the task, hampers result generalization. In the last decade, the study of functional connectivity at rest, using resting-state fMRI (rsfMRI), has gained interest to probe the sensorimotor system in CP. In this context, initial studies indicated that compared to healthy controls, functional connectivity of the sensorimotor network in the CP population seems more diffuse and widespread, leading to a potential reduced specificity and lower network efficiency (Burton, Dixit, Litkowski, & Wingert, 2009; J. D. Lee et al., 2011; Papadelis et al., 2014; Saunders, Carlson, Cortese, Goodyear, & Kirton, 2018). Moreover, the lateralization of the sensorimotor resting network toward the dominant hemisphere has been suggested to predict unimanual treatment response in uCP, highlighting its potential use as a biomarker for guiding clinical decision making (Manning et al., 2015; Rocca et al., 2013). Despite these interesting novel insights, previous studies have included small sample sizes (from 3 to 18 participants) with a rather heterogeneous population (different types of CP, i.e. unilateral and bilateral; different types of lesions, i.e. predominantly white matter versus grey matter). Furthermore, none of the prior studies have investigated the potential influence of different CST wiring patterns on functional connectivity of the sensorimotor network. Unravelling the potential relationships between aberrant functional connectivity, structural reorganization of the CST, and UL motor function might help to better understand the underlying mechanisms of sensorimotor dysfunction in uCP.

Given the lack of sufficient knowledge on the functional connectivity of the sensorimotor network in uCP compared to a large cohort of healthy controls, this study aims to investigate the occurrence of deviant functional connectivity of the sensorimotor network in a homogenous sample of 31 individuals with uCP due to white matter injury (i.e. periventricular leukomalacia or intraventricular haemorrhage) versus 60 healthy controls. Secondly, as CST wiring patterns have been put forward as one of the main factors influencing UL function, we specifically aimed to explore whether functional connectivity differs between different CST wiring groups (i.e., ipsilateral, contralateral, bilateral projections), and third, we explored the extent to which variations in functional connectivity and the type of CST wiring are predictive of UL function.

The following working hypotheses were tested in this study:

1. White matter lesions provoke deviant intra- and interhemispheric connectivity in the sensorimotor network (in dominant and non-dominant hemisphere), as compared to typically developing (TD) children.
2. The underlying CST wiring pattern alters sensorimotor functional connectivity in the uCP group, whereby alterations are more pronounced in the ipsilateral and bilateral groups.
3. The sensorimotor network in the uCP group is more widespread than in controls, and this differs according to the CST wiring pattern.
4. Upper limb motor deficits are related to sensorimotor functional connectivity measures and the combination of the underlying CST wiring and the connectivity measures will better explain the variability in motor deficits in uCP due to white matter lesions.

## 2. Materials and Methods

### 2.1. Participants

#### 2.1.1. Unilateral CP cohort

Thirty-one children, adolescents and young adults with uCP with a periventricular white matter lesion (PV lesion) were prospectively recruited via the CP reference center of the University Hospitals Leuven between 2014 and 2017. They were excluded if they had (1) botulinum toxin injections in the UL six months prior to the evaluation, (2) UL surgery two years prior to the assessment and/or (3) a comorbidity with other neurological or genetic disorders. All participants underwent a Magnetic Resonance Imaging (MRI) and Transcranial Magnetic Stimulation (TMS) session. According to the declaration of Helsinki, all participants assented to partake, signed the informed consent if >12 years old, and parents of participants <18 years old additionally signed the informed consent. This study was approved by the Ethics Committee Research UZ/KU Leuven (S55555 and S56513).

#### 2.1.2. Typically developing cohort

Sixty age-matched TD individuals were retrospectively selected from three sources. First, we screened the Autism Brain Imaging Data Exchange (ABIDE) database (Di Martino et al., 2014) (http://fcon_1000.proiects.nitrc.org/indi/abide/) and selected the Leuven 1 and Leuven 2 samples, due to the identical rsfMRI scanning procedure. Second, two ongoing studies at the KU Leuven recruited TD individuals for later comparison with clinical population, with identical scanning procedure (approved by the Ethics Committee Research UZ/KU Leuven (S25470 and S54757)). Finally, 26 were selected from ABIDE, 23 from study S25470, and 11 from study S54757.

### 2.2. MRI session

#### Data acquisition

Prior to the MRI, young children (were familiarized to the scanner situation in a playful manner during a training session using scan-related tasks that have been described elsewhere (Theys, Wouters, & Ghesquière, 2014). All participants (uCP and TD cohorts) underwent a single MR session in the same scanner machine acquired with a 3T system (Achieva, Philips Medical Systems, Best, The Netherlands) and equipped with a 32 channels coil in the University Hospitals Leuven (campus Gasthuisberg). Cushions were used to fix participants’ head in the coil to prevent motion artefacts.

Structural images were acquired using a 3D magnetization prepared rapid gradient echo (MPRAGE) [TR = 9.7ms, TE = 4.6ms, FOV = 250×250×192mm, voxel size = 0.98×0.98×1.2mm, acquisition time = 6 minutes]. Structural scans were inspected by a paediatric neurologist (EO) to ensure that only children with PV lesions were included in the analysis.

RsfMRI images were acquired using a T2*-weighted gradient-echo planar imaging (GE-EPI) [30 axial slices, slice thickness = 4 mm; no gap; TR = 1.7 s; TE = 33 ms; matrix size = 64×62; field of view = 230×230×120 mm; voxel size = 3.5×3.5×3.5 mm, flip angle = 90°; number of functional volumes = 250; acquisition time = 7 min]. Before the 250 volumes, four dummy volumes were acquired to stabilize the MR signal. Participants were instructed to lay still, not fall asleep and to think about nothing in particular.

#### Imaging pre-processing

Image pre-processing was conducted in SPM12 (www.fil.ion.ucl.ac.uk/spm). First, the structural images were registered to the T1 MNI template before the New Segmentation toolbox was used to segment the data into grey matter (GM), white matter (WM), and cerebro-spinal fluid (CSF) images. Next, functional images were co-registered to the individual structural images, realigned, and normalized to MNI space (resampled to 3×3×3 mm). After normalization, we flipped the structural and functional images of those with right-sided lesioned (in the uCP group) and left hemisphere dominance (i.e. right-handed participants in the TD cohort), so that the non-dominant and dominant hemispheres are on the same side. Throughout this manuscript, we use common terminology for both cohorts: dominant and non-dominant hemisphere, which corresponds to non-lesioned and lesioned hemisphere, respectively, in the uCP cohort. The CONN toolbox (www.nitrc.org/proiects/conn, RRID:SCR_009550) (Whitfield-Gabrieli & Nieto-Castanon, 2012) was used for denoising and the final connectivity analyses. Head motion was modelled to remove residual head motion, including 6 regressors that originated from the realignment and their derivatives, along with the first 5 principal component time series extracted from the WM and CSF masks (Chai, Castañán, Öngür, & Whitfield-Gabrieli, 2012). Lastly, spike-regression and linear detrending (Pruim, Mennes, Buitelaar, & Beckmann, 2015) were also applied before filtering the data in the band 0.01-0.15 Hz. Given the potential confounding effects of micro-movements on resting-state functional connectivity (Power, Barnes, Snyder, Schlaggar, & Petersen, 2012; Van Dijk, Sabuncu, & Buckner, 2012), all analyses were performed on ‘scrubbed’ data, i.e. eliminating those frames displaying frame-wise displacement (FD) exceeding 0.5 mm or frame-wise changes in brain image intensity exceeding 0.5 Δ%BOLD. Participants with a mean motion higher than FD>0.8 mm were not included in the final analysis (n=5 in uCP cohort, none in TD cohort).

#### Functional connectivity analyses

Functional connectivity analyses within the sensorimotor network were performed to explore potential alterations in the uCP group compared to controls. More specifically, connectivity was explored from bilateral primary motor cortex (M1) to a distributed network of sensorimotor regions including bilateral primary sensory cortex (S1), bilateral dorsal and ventral premotor cortex (PMd, PMv); and the supplementary motor cortex (SMA). For each of these regions, spherical regions of interest (ROI) with a radius of 6 mm were centred around MNI coordinates based on a recent meta-analysis investigating the three-dimensional location and boundaries of motor and premotor cortices (Figure 1) (Mayka, Corcos, Leurgans, & Vaillancourt, 2006). Note that a single midline ROI was adopted to represent the SMA proper region, resulting in a total of 9 ROIs. Further, since the ROI volume of S1 showed a slight overlap with M1 (42 voxels, i.e. 4.7% of the ROI volume), we attributed the overlapping voxels to the M1 volume (and therefore removed these voxels from the S1 volume). The MNI coordinates used for each ROI are reported in Supporting Information (Table S1).

**Figure 1.**
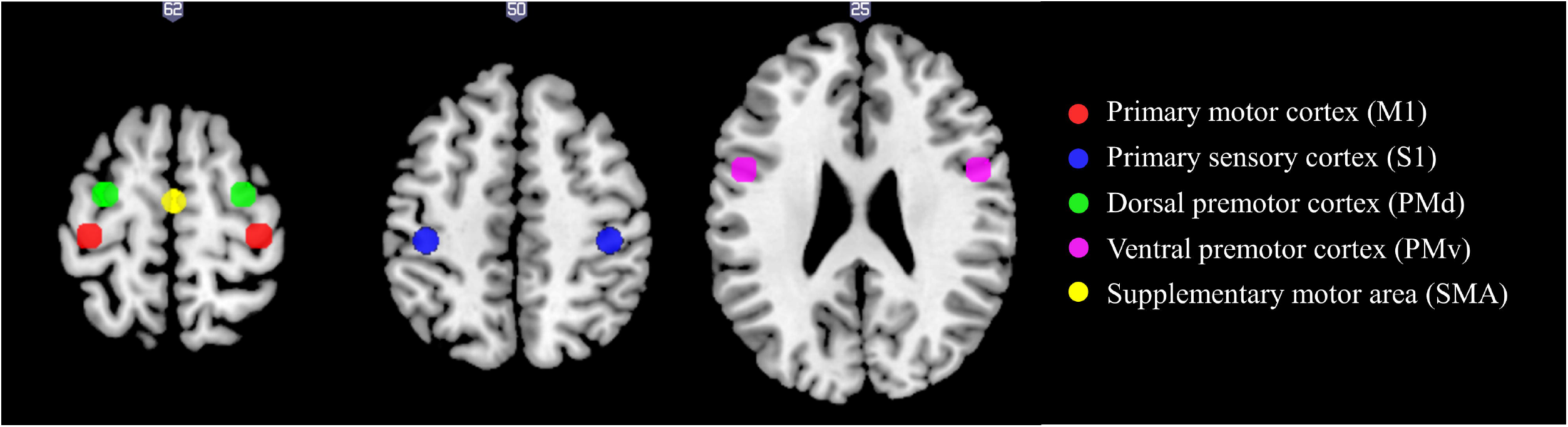
Regions of interest of the sensorimotor network included in the functional connectivity analyses in MNI space.

For each participant, we extracted the mean time-series of each ROI, calculated bivariate correlations between pairs of ROIs, and transformed the correlation coefficient to z-scores with the Fisher’s transformation. The connectivity measures included (i) ***intra***hemispheric functional connectivity of M1 with the other ROIs (S1, PMv, PMd) within each hemisphere (separate analyses for the non-dominant and dominant hemisphere); (ii) ***inter***hemispheric functional connectivity from M1 in the non-dominant hemisphere to the other ROIs of the dominant hemisphere (i.e. S1, PMd, PMv, and SMA) and vice versa; and (iii) ***inter***hemispheric functional connectivity between M1-M1.

Further, to investigate differences in intrahemispheric connectivity imbalance, we calculated the laterality index of the mean connectivity of all ROI pairs within one hemisphere according to the following formula (Seghier, 2008):

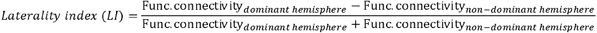

where a value closer to −1 would indicate complete laterality towards the non-dominant hemisphere, a value closer to +1 would indicate complete laterality toward the dominant hemisphere, and a value closer to 0 would indicate a balanced laterality (similar connectivity between hemispheres).

The primary motor network has been shown to be more diffuse and widespread in uCP, compared to controls (Vandermeeren, Davare, Duque, & Olivier, 2009). To explore this possibility in the current sample, we performed a secondary analysis, i.e. an exploratory seed-to-voxel based functional connectivity analysis, to identify remote connectivity of bilateral M1 to other brain regions not included in the ROI-ROI approach.

### 2.3. Transcranial Magnetic Stimulation (TMS)

To identify the CST wiring pattern in the uCP cohort (contralateral, ipsilateral, or bilateral), we conducted single-pulse TMS with a MagStim 200 stimulator (Magstim Ltd, Whitland, Wales, UK) equipped with a focal 70mm figure-eight coil and a Bagnoli electromyography (EMG) system with surface electrodes (Delsys Inc, Natick, MA, USA) attached to the adductor and opponens pollicis brevis muscles of both hands. A detailed description of the stimulation protocol can be found elsewhere (Simon-Martinez, Mailleux, et al., 2018). In short, hotspot and resting motor threshold were identified for each CST, by stimulating on the dominant hemisphere (i.e., identifying contralateral or potential ipsilateral projections), followed by the non-dominant hemisphere (i.e., identifying potential contralateral projections). Motor Evoked Potentials (MEPs) were bilaterally recorded to categorize all participants according to their underlying CST wiring pattern: contralateral, ipsilateral, or bilateral. All TMS measurements were conducted by two experienced physiotherapists (CSM and EJ).

### 2.4. Upper limb motor function evaluation

The Manual Ability Classification System (MACS) level was defined and reported for descriptive purposes (Eliasson et al., 2006). Grip strength, unimanual capacity and bimanual performance were evaluated in the uCP cohort. **Maximum grip strength** was assessed using the Jamar^®^ hydraulic hand dynamometer (Sammons Preston, Rolyan, Bolingbrook, IL, USA). The less-affected hand was measured first and the mean of three maximum contractions was calculated per hand. The ratio between hands was used for further analyses (grip strength ratio = less-affected hand/affected hand; i.e. a score closer to 1 indicates an adequate grip of the affected hand). **Hand dexterity** was assessed with the modified version of the Jebsen-Taylor hand function test (JTHFT) (Gordon, Charles, & Wolf, 2006; Taylor, Sand, & Jebsen, 1973). The time to perform every task was summed up and the ratio between hands was used for further analyses (JTHFT ratio = affected hand/less-affected hand; i.e. a score closer to 1 indicates an adequate dexterity of the affected hand). The Assisting Hand Assessment (AHA) was used to reliably measure **bimanual performance**, evaluating how effectively the affected hand is used in bimanual activities (Holmefur, Aarts, Hoare, & Krumlinde-Sundholm, 2009; Krumlinde-Sundholm & Eliasson, 2003; Krumlinde-Sundholm, Holmefur, Kottorp, & Eliasson, 2007). Given the age range of the participants, the School Kids AHA and the Ad-AHA were administrated (Louwers, Beelen, Holmefur, & Krumlinde-Sundholm, 2016). The AHA was scored by certified raters, using the 5.0 version, resulting in a final score between 0-100 AHA units. UL function was evaluated by experienced physiotherapists at the Clinical Motion Analysis Laboratory of the University Hospitals Leuven (campus Pellenberg, Belgium).

### 2.5. Statistical analyses

All behavioural data were checked for normality with the Shapiro-Wilk test and the histograms were inspected. Mean and standard deviation were reported for normally distributed data. If a non-normally distribution was found, a transformation was applied to allow parametric statistics.

First, we explored group differences in functional connectivity of the sensorimotor network between the uCP and the control cohort (**hypothesis #1**). Next, we investigated the impact of the CST wiring pattern by comparing sensorimotor functional connectivity of each CST wiring pattern with the control cohort and between wiring groups (**hypothesis #2**). For the first two hypotheses, we investigated group differences among the functional connectivity measures derived from the ROI-ROI approach at following levels: (1) ***intra***hemispheric functional connectivity of M1 within the ***non-dominant*** hemisphere in CP (non-dominant hemisphere in the control group), (2) ***intra***hemispheric functional connectivity of M1 within the ***dominant*** hemisphere, (3) ***inter***hemispheric functional connectivity between ***non-dominant M1*** to the ROIs on the dominant side; (4) ***inter***hemispheric functional connectivity between ***dominant M1*** to ROIs on the non-dominant side; and finally (5) ***inter***hemispheric functional connectivity between M1s. For each functional connectivity level, a repeated measures ANOVA model was conducted with the between-subject factor ‘group’ and the within-subject factor ‘connection’ (connectivity from M1 to the other ROIs) (Figure 2). The between-groups term was first entered to identify differences between uCP and TD individuals and secondly between TD individuals and each of the three uCP CST groups (contralateral, bilateral, and ipsilateral). Significant group*connection interactions were followed by post-hoc univariate ANOVAs for each ROI pair. If no interaction was found, between-group differences were reported. For the four-group comparison (hypothesis #2), post-hoc analyses were conducted if the main effect was significant and corrected for multiple comparison using Tukey’s HSD test. Lastly, differences in the laterality index were assessed with an ANOVA between groups (TD vs. uCP and TD vs. each CST wiring group).

**Figure 2.**
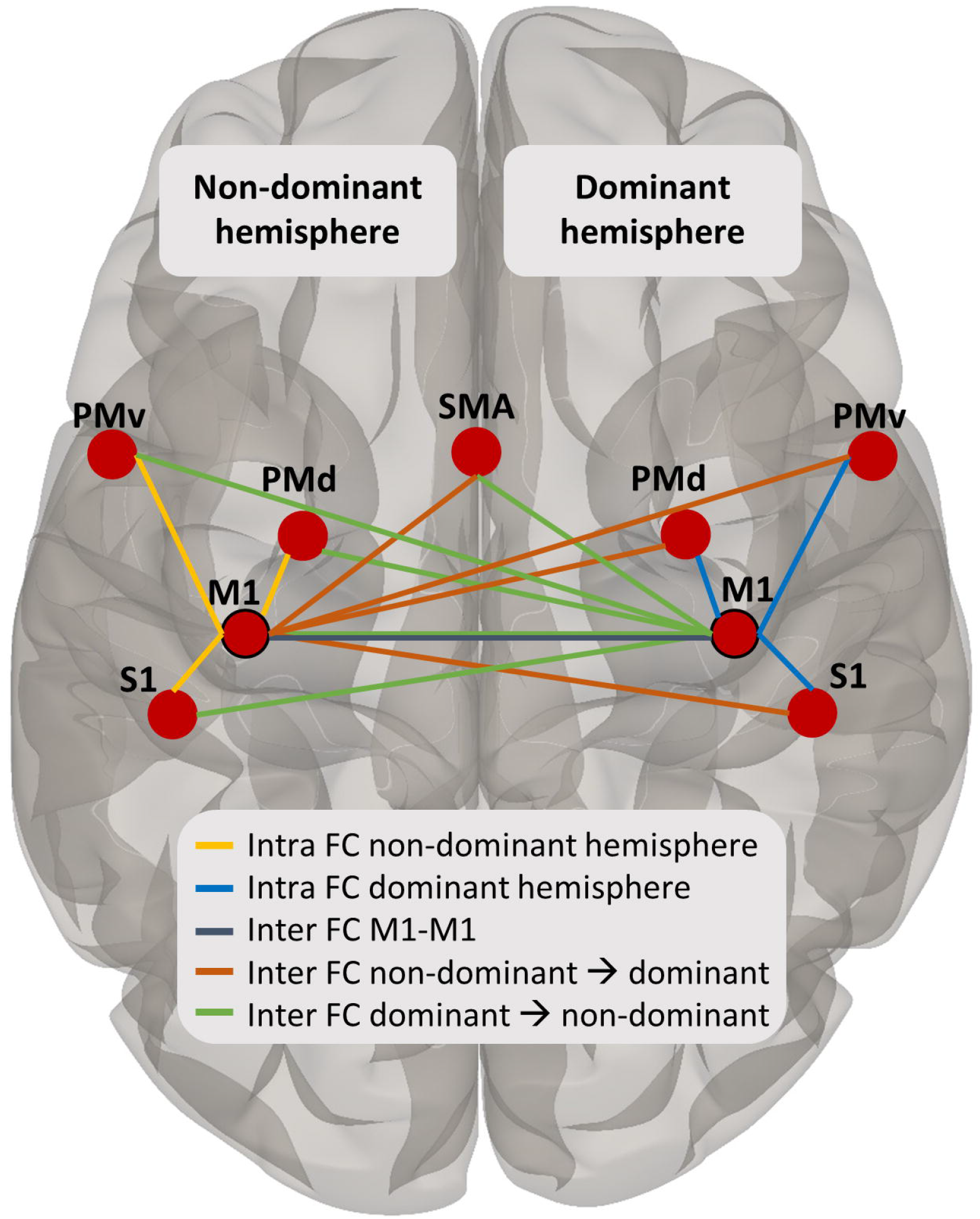
Overview of the ROIs included in the ROI-ROI analysis and the levels of functional connectivity tested in the statistical models: intrahemispheric functional connectivity in the non-dominant (in yellow) and dominant hemisphere (in blue), interhemispheric functional connectivity between M1-M1 (in dark grey), and interhemispheric functional connectivity from the non-dominant M1 to the dominant (in orange) and vice versa (in green).

Next to the ROI-ROI approach, exploratory seed-to-voxel functional connectivity analyses were also conducted. With this analysis, we aimed to identify differences in remote connectivity between each M1 (seeds) and other brain regions (**hypothesis #3**). We used a voxel-wise threshold p<0.001, and a cluster level p<0.05 to control the false discovery rate (FDR) (Benjamini & Hochberg, 1995; Genovese, Lazar, & Nichols, 2002), as implemented in SPM12 and the CONN toolbox (K. J. Friston, Ashburner, Kiebel, Nichols, & Penny, 2007; Whitfield-Gabrieli & Nieto-Castanon, 2012). The z-maps of each group (first TD vs. uCP, and then TD vs. each CST wiring group) were calculated and compared with an ANOVA contrast, to explore remote connectivity between uCP and TD and between each of the uCP CST wiring patterns and TD.

Lastly, correlation analyses were performed to evaluate whether the functional connectivity measures were related to motor deficits in the uCP group using Pearson’s r coefficients (**hypothesis #4**). Correlation coefficients <0.30 were considered little or no correlation, 0.30–0.50 low, 0.50–0.70 moderate, 0.70–0.90 high, and>0.90 very high (Hinkle & Wiersma, 1998). To evaluate the combined predictive value of the functional connectivity measures and the CST wiring pattern, we additionally conducted a multiple regression analysis. The functional connectivity measures entered in the model were selected based on the distinct connectivity pattern shown by the uCP group in the previous comparisons (TD vs. uCP group and TD vs. each CST wiring pattern). Interaction terms between the CST wiring patterns and the functional connectivity measures were also entered in the model, which was fitted using the backward selection method.

The alpha-level was set at 0.05 for interaction term, main effects, and correlation/regression analyses. Statistical analyses were performed using SPSS (Windows version 25.0, IBM Corp., Armonk, NY).

## 3. Results

After exclusion of high motion participants (mean FD>0.8, n=5), the final uCP sample included 26 individuals (15 girls; 12 right-sided uCP; 9 with MACS I, 11 with MACS II and 6 with MACS III) and 60 individuals in the TD cohort (14 girls; 54 right-handed). Age did not differ between groups (uCP cohort (X(SD)) = 12.87 (4.45); TD cohort (X(SD)) = 14.54 (4.80); p=0.10)). In the uCP cohort, we identified 9 individuals with a contralateral CST wiring, 6 with a bilateral, and 9 with an ipsilateral (two participants declined to participate in the TMS session; demographic data Supporting Information Table S2). Tables 1 and 2 summarize the functional connectivity measures in each group as derived from the ROI-ROI approach.

### 3.1. TD vs. uCP group differences in functional connectivity (hypothesis #1)

#### Intrahemispheric functional connectivity

Within the ***non-dominant hemisphere***, rmANOVA analyses with the between-subject factor ‘group’ (uCP, TD) and the within-subjects factor ‘intrahemispheric M1-connectivity’ (M1-PMd, M1-PMv, M1-S1) showed no differences in M1 intrahemispheric functional connectivity between groups (main effect of group, p=0.25) and also no significant interaction effect group*connection (F (2, 83) = 0.77, p=0.47, Wilks’ Lambda=0.98) (Table 1, Figure 3A). Within the ***dominant hemisphere***, rmANOVA analyses with the between-subject factor ‘group’ (uCP, TD) and the within-subjects factor ‘intrahemispheric M1-connectivity’ (M1-PMd, M1-PMv, M1-S1) showed no differences between groups (p=0.10) and no interaction effect (F (2, 83) = 1.13, p=0.32, Wilks’ Lambda=0.97) (Table 1, Figure 3B).

**Table 1.** Descriptive (mean (95% CI)) and comparative statistics of each ROI pair in each cohort for the uCP vs. TD comparison.

|  | TD cohort<br>(n=60) | uCP cohort<br>(n=26) | Wilk's Lambda (p)<br>group*connection | F (p)<br>main effect group |
| --- | --- | --- | --- | --- |
| **Intra functional connectivity non-dominant hemisphere** |  |  | 0.98 (0.47) | 1.33 (0.25) |
| M1-PMd | 0.33 (0.05) | 0.33 (0.09) |  |  |
| M1-PMv | 0.03 (0.03) | 0.09 (0.05) |  |  |
| M1-S1 | 0.69 (0.06) | 0.73 (0.10) |  |  |
| **Intra functional connectivity dominant hemisphere** |  |  | 0.97 (0.32) | 2.73 (0.10) |
| M1-PMd | 0.30 (0.06) | 0.39 (0.11) |  |  |
| M1-PMv | 0.03 (0.03) | 0.14 (0.07) |  |  |
| M1-S1 | 0.87 (0.07) | 0.85 (0.14) |  |  |
| **Inter functional connectivity Les → NonLes** |  |  | 0.92 (0.07) | 0.12 (0.73) |
| M1-PMd | 0.19 (0.05) | 0.27 (0.08) |  |  |
| M1-PMv | 0.05 (0.03) | 0.09 (0.06) |  |  |
| M1-S1 | 0.40 (0.06) | 0.34 (0.07) |  |  |
| M1-SMA | 0.32 (0.06) | 0.30 (0.09) |  |  |
| **Inter functional connectivity NonLes → Les** |  |  | 0.94 (0.14) | 0.04 (0.85) |
| M1-PMd | 0.22 (0.04) | 0.25 (0.07) |  |  |
| M1-PMv | 0.01 (0.03) | 0.09 (0.08) |  |  |
| M1-S1 | 0.38 (0.06) | 0.35 (0.07) |  |  |
| M1-SMA | 0.29 (0.06) | 0.23 (0.07) |  |  |
| M1-M1 | 0.47 (0.07) | 0.47 (0.09) |  | 0.00 (0.99) |

TD, typically developing; uCP, unilateral cerebral palsy; CST, corticospinal tract; M1, primary motor cortex; PMd, dorsal stream of the premotor cortex; PMv, ventral stream of the premotor cortex; S1, primary sensory cortex; SMA, supplementary motor area.

**Table 2.** Descriptive (mean (95% CI)) and comparative statistics of each ROI pair in each cohort for the TD vs. CST wiring group comparison.

|  | TD cohort<br>(n=60) | Contralateral<br>CST (n=9) | Bilateral<br>CST (n=6) | Ipsilateral<br>CST (n=9) | Wilk's Lambda (p)<br>group*connection | F (p)<br>main effect<br>group |
| --- | --- | --- | --- | --- | --- | --- |
| **Intra functional connectivity non-dominant hemisphere** |  |  |  |  | 0.95 (0.63) | 0.93 (0.43) |
| M1-PMd | 0.33 (0.05) | 0.35 (0.15) | 0.35 (0.26) | 0.24 (0.08) |  |  |
| M1-PMv | 0.03 (0.03) | 0.12 (0.11) | 0.03 (0.04) | 0.09 (0.10) |  |  |
| M1-S1 | 0.69 (0.06) | 0.77 (0.20) | 0.70 (0.16) | 0.68 (0.17) |  |  |
| **Intra functional connectivity dominant hemisphere** |  |  |  |  | **0.79 (0.005)\*** | - |
| M1-PMd | 0.30 (0.06) | 0.56 (0.21) | 0.42 (0.22) | 0.23 (0.10) | 4.16 (0.009)<sup>‡§</sup> |  |
| M1-PMv | 0.03 (0.03) | 0.09 (0.13) | 0.18 (0.13) | 0.15 (0.11) | 3.59 (0.02) |  |
| M1-S1 | 0.87 (0.07) | 0.77 (0.17) | 0.99 (0.43) | 0.77 (0.19) | 0.88 (0.45) |  |
| **Inter functional connectivity Les → NonLes** |  |  |  |  | 0.89 (0.37) | 0.47 (0.70) |
| M1-PMd | 0.19 (0.05) | 0.27 (0.13) | 0.28 (0.19) | 0.24 (0.16) |  |  |
| M1-PMv | 0.05 (0.03) | 0.06 (0.09) | 0.10 (0.07) | 0.10 (0.13) |  |  |
| M1-S1 | 0.40 (0.06) | 0.28 (0.11) | 0.34 (0.11) | 0.39 (0.12) |  |  |
| M1-SMA | 0.32 (0.06) | 0.21 (0.13) | 0.29 (0.16) | 0.35 (0.18) |  |  |
| **Inter functional connectivity NonLes → Les** |  |  |  |  | 0.87 (0.29) | 0.03 (0.99) |
| M1-PMd | 0.22 (0.04) | 0.28 (0.14) | 0.28 (0.15) | 0.22 (0.13) |  |  |
| M1-PMv | 0.01 (0.03) | 0.07 (0.18) | 0.03 (0.04) | 0.14 (0.14) |  |  |
| M1-S1 | 0.38 (0.06) | 0.40 (0.16) | 0.34 (0.10) | 0.32 (0.11) |  |  |
| M1-SMA | 0.29 (0.06) | 0.17 (0.10) | 0.30 (0.14) | 0.21 (0.16) |  |  |
| M1-M1 | 0.47 (0.07) | 0.43 (0.13) | 0.47 (0.13) | 0.50 (0.11) |  | 0.10 (0.96) |

TD, typically developing; uCP, unilateral cerebral palsy; CST, corticospinal tract; M1, primary motor cortex; PMd, dorsal stream of the premotor cortex; PMv, ventral stream of the premotor cortex; S1, primary sensory cortex; SMA, supplementary motor area. \*Statistically significant (p<0.05).
<sup>‡</sup>Contralateral CST vs. Ipsilateral CST; <sup>§</sup>Contralateral CST vs. TD group.

**Figure 3.**
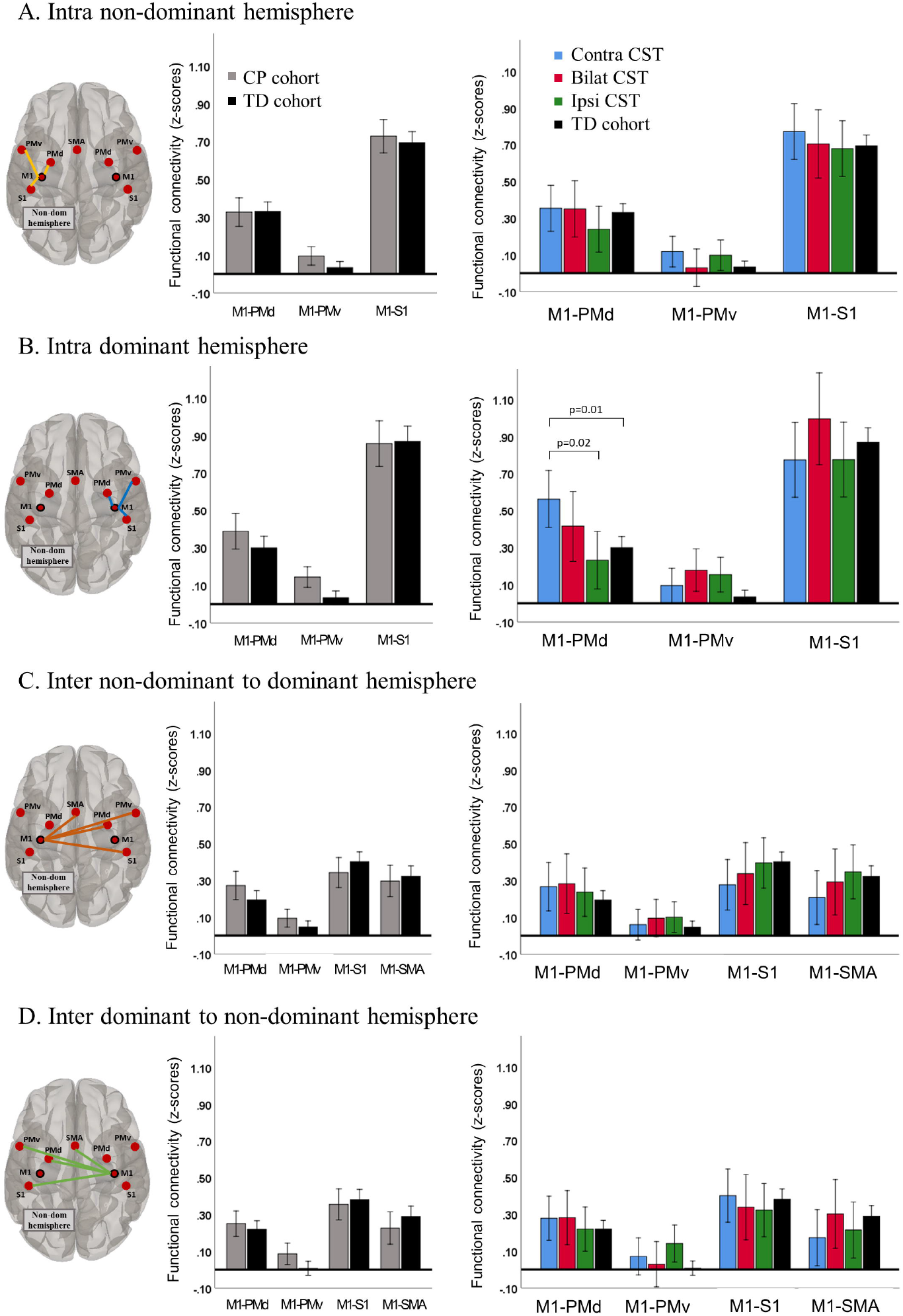
Functional connectivity patterns between uCP and TD cohort (left panel) and each CST wiring group and the TD cohort (right panel) tested at four levels: Intrahemispheric connectivity in the non-dominant hemisphere (A), and in the dominant hemisphere (B); and interhemispheric connectivity from the non-dominant M1 to the dominant-sided ROIs (C), and from the dominant M1 to the non-dominant-sided ROIs (D). Bars illustrate mean and 95% confidence interval for each group.

#### Imbalance between intrahemispheric functional connectivity

Figure 4a shows the laterality indices in each group. We found no differences in intrahemispheric imbalance between the uCP and TD cohorts (p=0.38).

**Figure 4.**
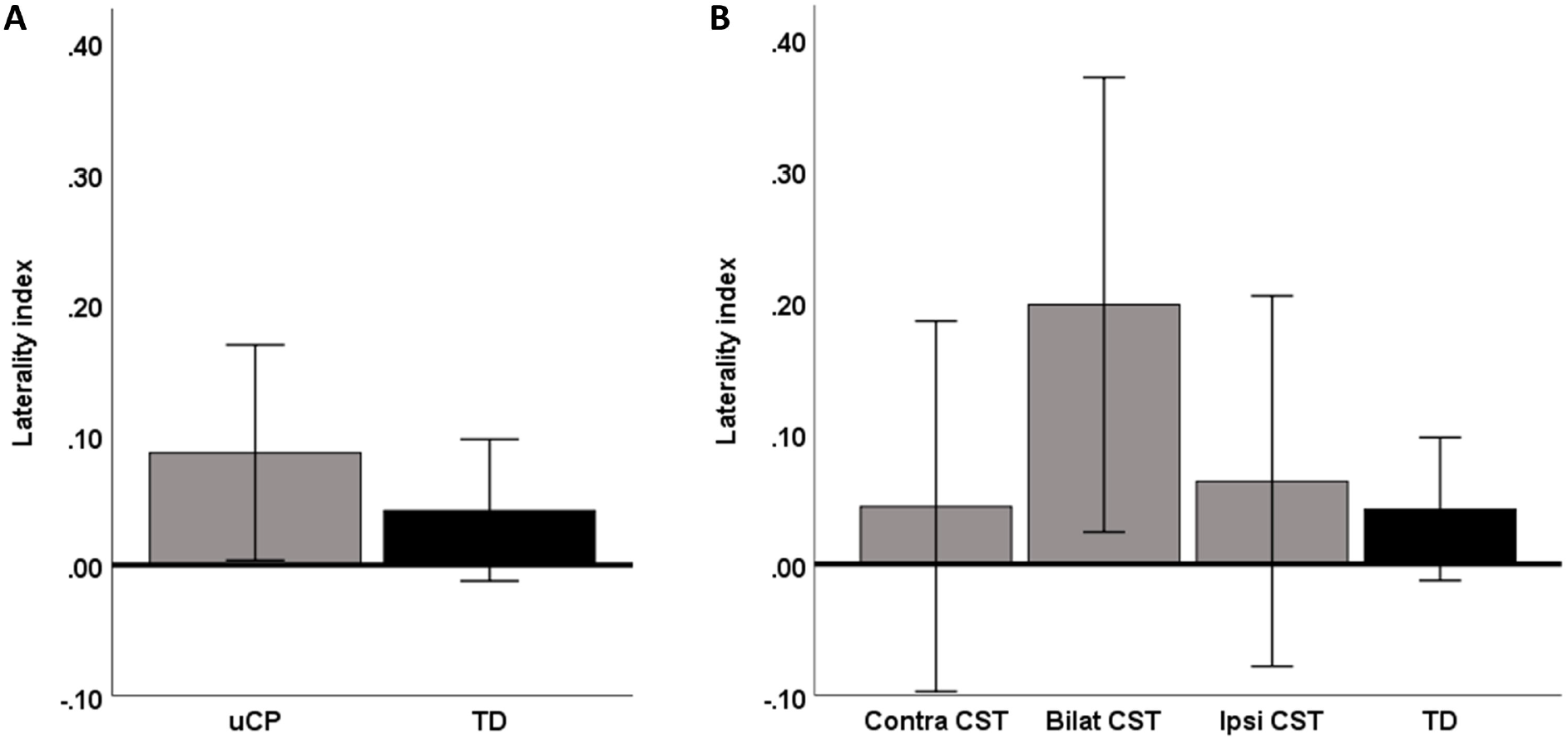
Laterality indices of the intrahemispheric connectivity in each group, highlighting the balance in the TD cohort, whose index is close to 0. (A) Comparison between TD and uCP group; (B) comparison between TD and CST wiring groups. Bars indicate the group mean and error bars indicate the 95% confidence interval.

#### Interhemispheric functional connectivity

For interhemispheric connectivity between ***non-dominant M1*** and sensorimotor ROIs in the contralateral, dominant hemisphere, rmANOVA analyses with the between-subject factor ‘group’ (uCP, TD) and the within-subjects factor ‘interhemispheric M1-connectivity’ showed a trend toward a ‘group*connection’ interaction (F (3, 82) = 2.45, p=0.07, Wilks’ Lambda=0.92), indicating a different pattern between ROI pairs in each group (Table 1, Figure 3C). The main effect of group was not significant (p=0.73). Secondly, the interhemispheric connectivity pattern between ***dominant M1*** and sensorimotor ROIs in the non-dominant hemisphere did not show an interaction effect between group (uCP, TD) and connectivity measures (F (3, 82) = 1.86, p=0.14, Wilks’ Lambda=0.94), nor a group effect (p=0.85) (Table 1, Figure 3D). Lastly, the interhemispheric connectivity between ***M1-M1*** was not different between the uCP and TD cohort (F (1, 84) < 0.01, p=0.99) (Table 1).

In conlusion, we do not find evidence in support of hypothesis 1, suggesting that functional connectivity is not uCP-dependent.

### 3.2. TD vs. CST wiring group differences in functional connectivity (hypothesis #2)

#### Intrahemispheric functional connectivity

Similar to the first hypothesis, differences in terms of intrahemispheric functional connectivity ***within the non-dominant hemisphere*** between the TD group and each of the CST wiring groups were not significant (interaction group*connection, F (6, 158) = 0.73, p=0.63, Wilks’ Lambda = 0.95; main effect of group, p=0.43). The connectivity pattern in intrahemispheric functional connectivity ***within the dominant hemisphere*** between the CST wiring groups and the TD cohort showed a significant group*connection interaction (F (6, 158) = 3.28, p=0.005, Wilks’ Lambda=0.79). The univariate results indicated that the group differences were mainly driven by differential connectivity between M1-PMd (p=0.009) and M1-PMv (p=0.017) (Figure 3B). Post-hoc analyses for the M1-PMd connectivity depicted higher connectivity in the contralateral compared to the ipsilateral CST group (p=0.018, Tukey HSD corrected) and the TD cohort (p=0.011, Tukey HSD corrected). Post-hoc analysis for M1-PMv showed that the connectivity was tentatively higher in the bilateral and ipsilateral CST groups compared to the TD cohort, although post-hoc analyses did not survive multiple comparison correction (both p=0.09, Tukey HSD corrected).

#### Imbalance between intrahemispheric functional connectivity

Figure 4b shows the laterality indices in each group. We found no differences between TD and each CST wiring group (p=0.40).

#### Interhemispheric functional connectivity

The interhemispheric functional connectivity from ***non-dominant M1*** did not differ when comparing the CST wiring groups and the TD group (no interaction effect (F (9, 190) = 1.09, p=0.37, Wilks’ Lambda=0.89); no group effect (p=0.70)). Similarly, from ***dominant M1***, the analysis comparing the CST wiring groups and the TD group showed no interaction effect (F (9, 190) = 1.21, p=0.29, Wilks’ Lambda=0.87), and no group effect (p=0.99). Lastly, the connectivity between ***M1-M1*** was not different between the CST wiring and the TD group (F (3, 80) = 0.10, p=0.96).

In summary, we find evidence in support of hypothesis 2, suggesting that intrahemispheric functional connectivity within the dominant hemisphere is CST-wiring dependent, specifically between the primary and premotor cortices.

### 3.3. Seed-to-voxel analysis exploring remote M1 functional connectivity (hypothesis #3)

Seed-to-voxel analyses were performed to explore group differences in remote functional connectivity from each M1 to all the other voxels in the brain. Figure 5 shows the connectivity pattern from each M1 in every group (TD, contralateral CST, ipsilateral CST, and bilateral CST).

**Figure 5.**
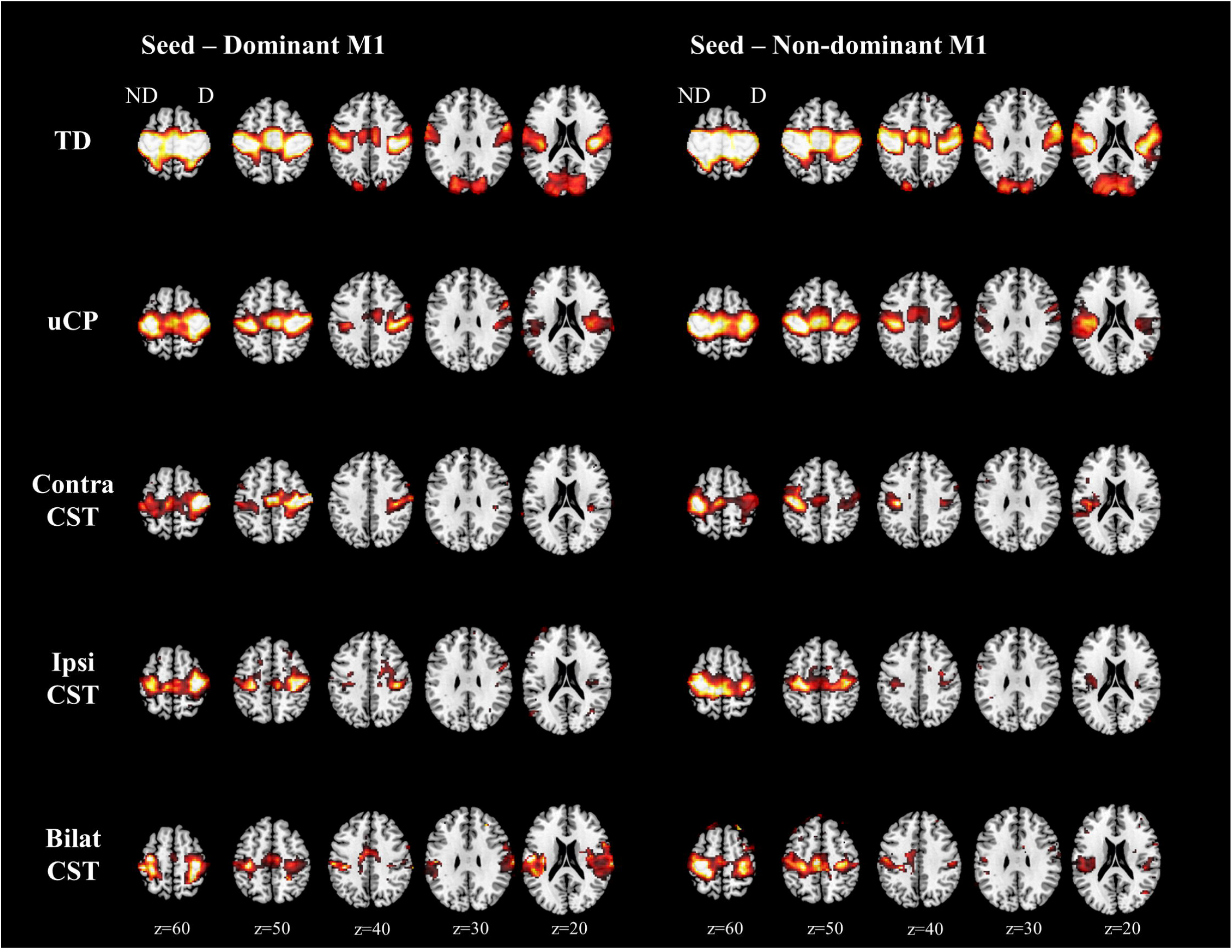
Functional connectivity from each M1 to all the other voxels in the brain. T-maps are thresholded to the one-sample t-test for the TD group (t=3.24). TD, typically developing; uCP, unilateral cerebral palsy; CST, corticospinal tract; ND, non-dominant hemisphere; D, dominant hemisphere.

First, we investigated differences between the TD group and uCP group (Table 3A). Similar to the ROI-ROI analyses, we found no group differences **from the non-dominant M1** with other sensorimotor areas, although we found higher functional connectivity between M1 and both occipital poles in the TD cohort, compared to the uCP group (non-dominant-side occipital pole (intrahemispheric functional connectivity), p-FDR corrected <0.001; dominant side occipital pole (interhemispheric functional connectivity), p-FDR corrected <0.001). In contrast, the uCP group showed higher functional connectivity between non-dominant M1 and the ipsilateral temporal pole and the insular cortex (p-FDR corrected =0.01) (Figure 6). **From the dominant M1**, no differences were identified between groups.

**Table 3A.** Differences in connectivity pattern from non-dominant M1 to other brain regions.

| Seed on non-dominant M1 |  |  |  |  |  |  |  |
| --- | --- | --- | --- | --- | --- | --- | --- |
|  | Cluster # | Location | Direction | Coordinates (x,y,z) | p-unc | p-FDR corrected | Cluster size |
| TD vs. uCP | 1 | Occipital pole (non-dominant side) | TD > uCP | -11, -97, +31 | <0.001 | <0.001 | 8684 |
|  | 2 | Occipital pole (dominant side) | TD > uCP | +27, -101, +16 | <0.001 | <0.001 | 8109 |
|  | 3 | Temporal pole, insular cortex (non-dominant side) | uCP > TD | -53, +16, -14 | 0.001 | 0.01 | 2804 |
| TD vs. CST wiring | 1 | Occipital pole (non-dominant side) | TD > contra<br>TD > ipsi<br>TD > bilat | -26, -95, +10 | <0.001 | <0.001 | 7273 |
|  | 2 | Occipital pole (dominant side) | TD > contra<br>TD > ipsi<br>TD > bilat<br>Ipsi > bilat | +10, -95, +28 | <0.001 | <0.001 | 7140 |

TD, typically developing; uCP, unilateral Cerebral Palsy; CST, corticospinal tract; M1, primary motor cortex; p-unc, p-uncorrected; FDR, False Discovery Rate.

**Table 3B.** Differences in connectivity pattern from dominant M1 to other brain regions.

| Seed on dominant M1 |  |  |  |  |  |  |  |
| --- | --- | --- | --- | --- | --- | --- | --- |
|  | Cluster # | Location | Direction | Coordinates (x,y,z) | p-unc | p-FDR corrected | Cluster size |
| TD vs. uCP | None |  |  |  |  |  |  |
| TD vs. CST wiring | 1 | Occipital pole (non-dominant side) | TD > contra<br>TD > bilat<br>Ipsi > bilat | -17, -95, +16 | <0.001 | <0.001 | 3585 |
|  | 2 | Occipital pole (dominant side) | TD > contra<br>TD > bilat<br>Ipsi > bilat | +19, -95, +16 | <0.001 | 0.002 | 6426 |
|  | 3 | Parietal operculum, supramarginal gyrus (dominant side) | Bilat > contra<br>Bilat > ipsi<br>Bilat > TD | +55, -29, +25 | <0.001 | 0.002 | 3483 |

TD, typically developing; uCP, unilateral Cerebral Palsy; CST, corticospinal tract; M1, primary motor cortex; p-unc, p-uncorrected; FDR, False Discovery Rate.

**Figure 6.**
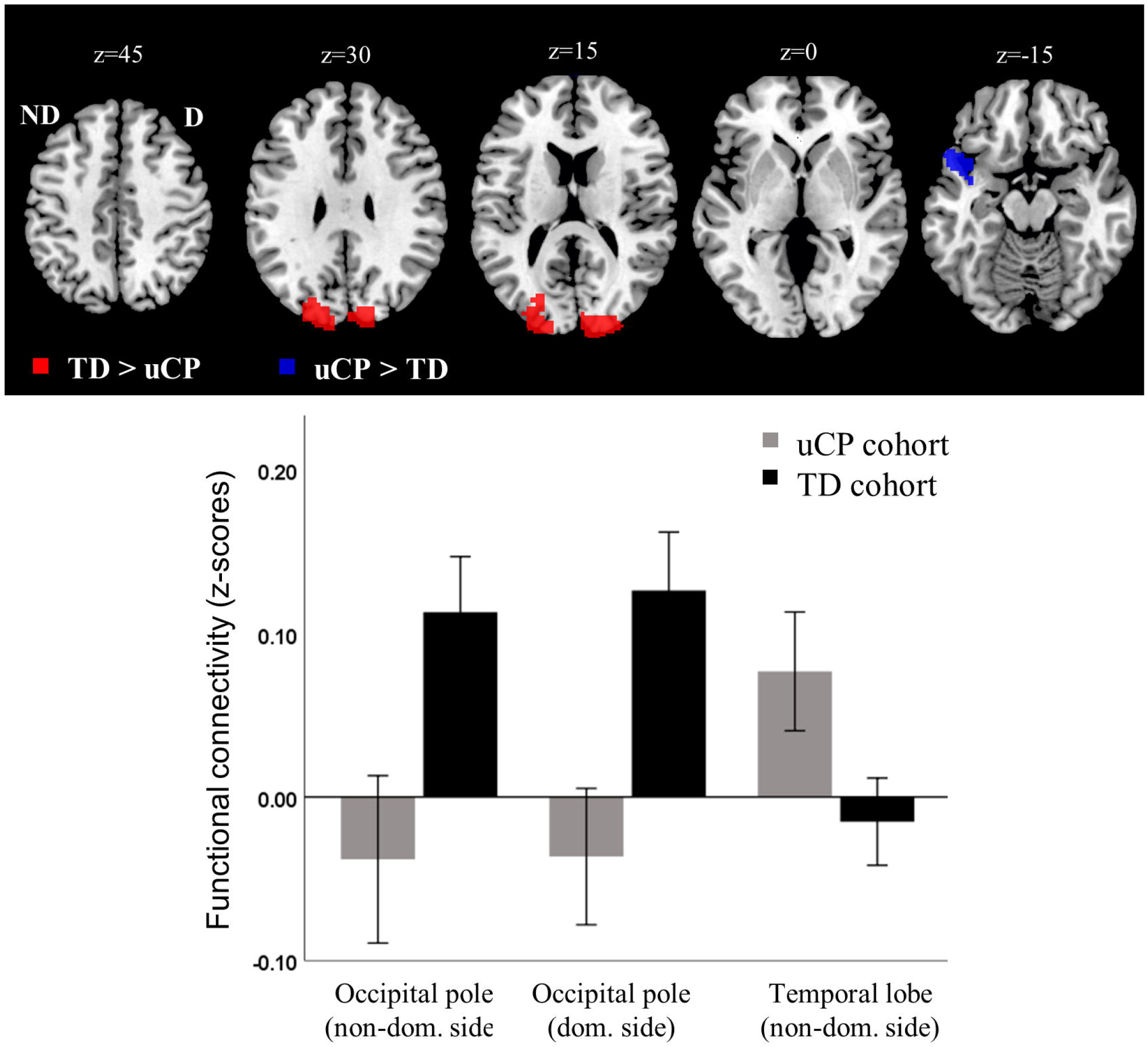
Increased uCP-dependent functional connectivity from non-dominant side M1 was identified in the temporal lobe, whereas lower connectivity was found in the occipital poles. uCP, unilateral cerebral palsy; TD, typically developing; ND/non-dom., non-dominant side; D/dom., dominant side.

Secondly, we explored differences between the TD cohort and each of the CST wiring groups (Table 3B). **From the non-dominant M1**, group differences were found in the non-dominant-side occipital pole (i.e. intrahemispheric functional connectivity; p-FDR corrected <0.001) and in the contralateral occipital pole (i.e. interhemispheric functional connectivity; p-FDR corrected <0.001). Post-hoc analysis indicated that the TD group had higher connectivity than any of the CST wiring groups (p-FDR corrected <0.05). Figure 7A shows the functional connectivity data of each group, illustrating the low connectivity in each CST wiring group, despite the group differences. From the dominant hemisphere group differences were also found in both occipital poles (dominant-side occipital pole, i.e. intrahemispheric functional connectivity; p-FDR corrected <0.01); non-dominant-side occipital pole, i.e. interhemispheric functional connectivity; p-FDR corrected <0.001). Post-hoc analyses indicated for both clusters a similar pattern: higher connectivity in the TD cohort compared to the contralateral and bilateral CST wiring groups (p-FDR corrected <0.05), and higher connectivity in the ipsilateral compared to the bilateral CST group (p-FDR corrected = 0.04). Interestingly, we also identified a cluster covering the ipsilateral supramarginal gyrus, and the parietal operculum (i.e. intrahemispheric functional connectivity; p-DFR corrected = 0.002). Post-hoc analysis indicated higher intrahemispheric functional connectivity in the bilateral CST group compared to the contralateral CST (p-FDR <0.001), the ipsilateral CST (p-FDR <0.001), and the TD group (p-FDR corrected <0.001) (Figure 7B).

**Figure 7.**
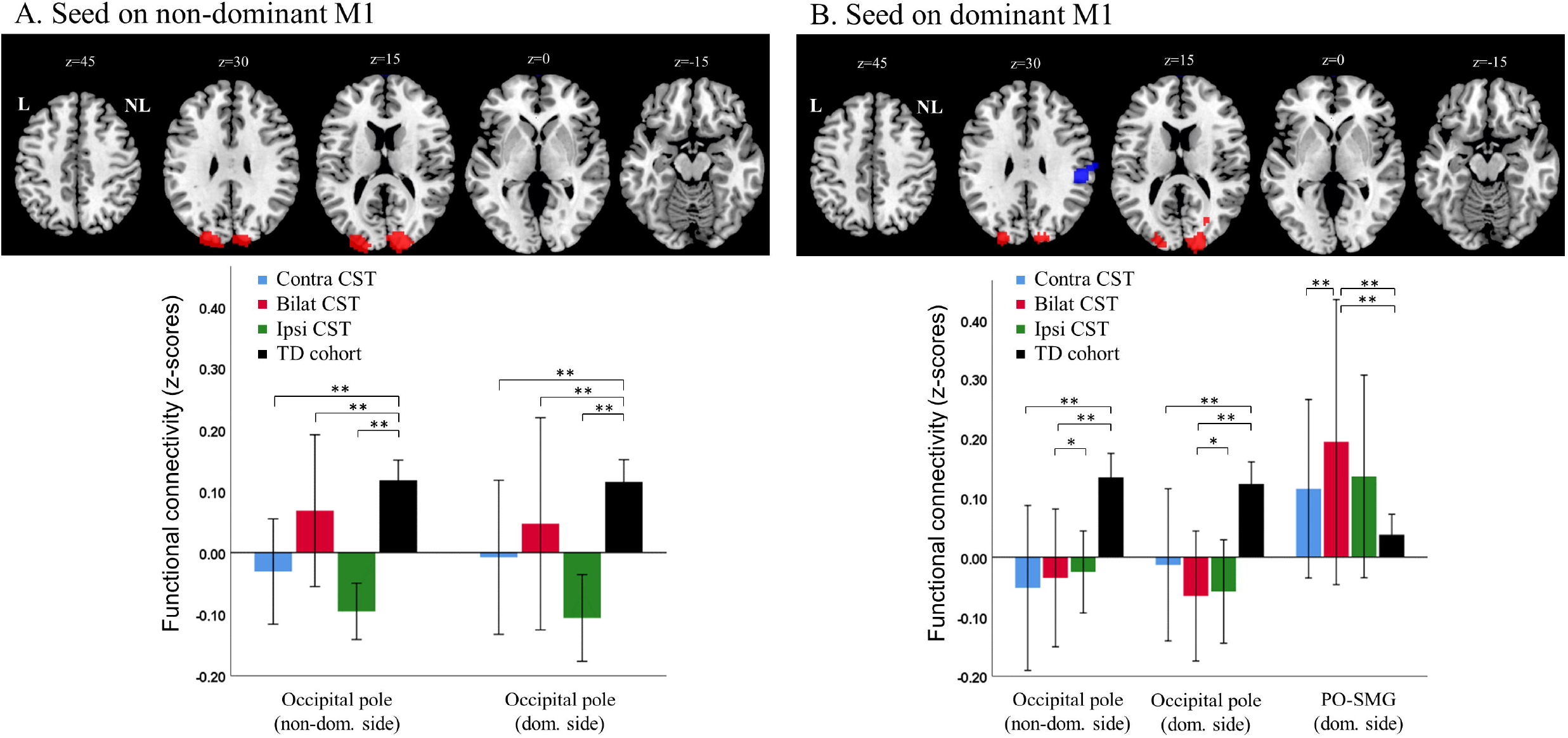
Functional connectivity CST wiring dependent from the non-dominant M1 (A) and from the dominant M1 (B). Seed-to-voxel analysis depicted differences in functional connectivity between both M1 and the occipital poles, as well as between dominant M1 and association areas within the same hemisphere. PO-SMG, parietal operculum and supramarginal gyrus; CST, corticospinal tract; TD, typically developing; Dom, dominant; M1, primary motor cortex.

In conclusion, we find evidence in support of hypothesis 3, suggesting that there exist a more widespread sensorimotor network, that is CST-wiring dependent, specifically with somatosensory association areas.

### 3.4. Influence of functional connectivity and CST wiring pattern on motor function (hypothesis #4)

#### Correlation analysis

For the uCP cohort, no to low correlations were found between functional connectivity measures and UL motor deficits. The interhemispheric functional connectivity showed low correlations (−0.28 to - 0.30) between non-dominant M1 and dominant SMA with bimanual performance and hand dexterity, although they did not reach significance. The interhemispheric functional connectivity between dominant M1 and contralateral S1 tended to correlate with grip strength (r=−0.36, p=0.08), hand dexterity (r=−0.39, p=0.05), and bimanual performance (r=0.35, p=0.09), whereby higher connectivity indicated better motor function (Supporting Information, Table S3).

#### Regression analysis

The regression analysis included the connectivity measures based on the ROI approach that were uCP- and CST-dependent (i.e., intrahemispheric connectivity in the dominant hemisphere: (i) M1-PMd, (ii) M1-PMv, and (iii) the cluster identified in the seed-to-voxel approach (dominant M1 to parietal operculum and supramarginal gyrus) to predict grip force, hand dexterity, and bimanual performance. The results showed that the measures of intrahemispheric connectivity of the dominant hemisphere were not able to predict UL motor function (grip strength R^2^ = 0.04, p=0.83; hand dexterity R^2^ = 0.15, p=0.32; and bimanual performance R^2^ = 0.05, p=0.80).

In a second step, the underlying type of CST wiring was included into the model as an interacting and main effect. The backward selection method only retained the type of CST in the model as able to predict grip strength (R^2^=0.33, p=0.02) and bimanual performance (R^2^=0.39, p=0.007). Interestingly, the connectivity derived from the cluster covering the somatosensory association areas (from dominant M1 to dominant-sided somatosensory association areas) tended to significantly contribute to hand dexterity, in combination with the underlying type of CST wiring (R^2^=0.37, CST wiring p=0.03, functional connectivity p=0.10), whereby higher connectivity in the dominant hemisphere and having an ipsilateral or bilateral CST wiring predicted poorer dexterity.

Briefly, we do not find evidence in support of hypothesis 4, indicating a small relationship between functional connectivity and UL motor function measures, which suggests that the underlying type of CST wiring remains the main predictor of motor function.

## 4. Discussion

In this study, we investigated differences in functional connectivity of the sensorimotor network based on rsfMRI, in a cohort of individuals with uCP with homogeneous brain damage (due to periventricular white matter injuries) and a large group of healthy age-matched controls. We included the type of CST wiring in the uCP group to explore functional connectivity differences between the CST wiring groups and examined the ability of these two measures (i.e. functional connectivity and CST wiring) to explain the underlying pathophysiology of UL motor problems. To do this, we chose an ROI-ROI approach to identify deviant connectivity patterns between core regions of the sensorimotor network, and a seed-to-voxel approach to elucidate whether aberrant functional connectivity may exist with other brain areas (i.e. remote connectivity due to compensation). Despite the lack of uCP-dependent aberrant connectivity compared to controls, as identified by the ROI-ROI approach, we found that the strength in the connectivity measures between M1 and the premotor cortices in the dominant hemisphere was dependent on the type of CST wiring. The seed-to-voxel approach also identified somatosensory association areas where the connectivity pattern was dependent on the type of CST wiring. Nevertheless, our results confirm that the CST wiring remains the main predictor of UL motor deficits, whereas functional connectivity seems to have little predictive value.

Our **first hypothesis** stated that white matter lesions in uCP would provoke deviant functional connectivity at the intra- and interhemispheric level, compared to controls, which cannot be fully rejected. The lack of differences in intrahemispheric connectivity within sensorimotor areas (in the ROI-ROI approach) in the non-dominant hemisphere was unexpected, as we hypothesized that the white matter lesion would reflect changes in this network. A recent study by Saunders et al. has also shown that the resting state motor network in children with such injuries highly resembles the motor network of TD peers (Saunders et al., 2018). However, they found differences in the laterality index between the two primary motor cortices, with grater asymmetry from the lesioned hemisphere, which we could not replicate. With our larger sample size, we can still observe that the functional connectivity network of the whole uCP sample due to periventricular white matter lesions very well resembles that of the TD cohort. Together with previous literature (Saunders et al., 2018), our results suggest that the brain in individuals with a periventricular white matter lesion may have higher plasticity potential to reorganize the motor network in such a way that it is not functionally altered. Other lesion types occurring around birth, i.e. ischemic arterial stroke, when grey matter is predominantly damaged, seem to show a more lateralized motor network (Saunders et al., 2018), potentially due to the grey matter loss.

Our **second hypothesis** stated that the functional connectivity is dependent on the type of CST wiring, which was confirmed by our results. In short, M1-PMd connectivity in the dominant hemisphere was higher in the contralateral CST group compared to the ipsilateral and the TD groups, whilst the M1-PMv connectivity was higher in the ipsilateral and bilateral CST groups compared to the TD group. Although both PMd and PMv have been shown to contribute to movement preparation and visuomotor transformation during grasping (Jeannerod, Arbib, Rizzolatti, & Sakata, 1995), these two areas seem to have a disparate role in controlling grasp function: PMd controls the reaching and the coupling between the grasping and lifting phases, whilst PMv mainly contributes to the grasping component (Davare, Andres, Cosnard, Thonnard, & Olivier, 2006). As individuals with a contralateral CST usually present with adequate motor function, the increased connectivity between M1-PMd may be a compensation for the finer features of grasping (i.e. the coupling between grasping and lifting), which necessitates from a synchrony between proximal and distal muscles, (Gupta et al., 2017; Simon-Martinez, Jaspers, et al., 2018; Zewdie et al., 2017). On the other hand, the increased connectivity between M1-PMv within the dominant hemisphere of the bilateral and ipsilateral CST wiring groups could suggest a prioritization of the grasping component, as the individuals with these types of CST wiring show poorer UL motor function (Simon-Martinez, Jaspers, et al., 2018; Staudt et al., 2002). Moreover, it is reasonable that the increased connectivity is present in the dominant hemisphere of the bilateral and ipsilateral CST wiring groups, as this hemisphere is the one with the main motor output, certainly in the ipsilateral group. To what extent an increased connectivity between M1-PMv within the dominant hemisphere has an impact on behaviour remains unknown, as we found only low correlations with UL motor function.

Regarding the interhemispheric functional connectivity in the uCP cohort within our second hypothesis, we did not find differences in interhemispheric functional connectivity between groups and this measure does not seem to be related to the underlying type of CST wiring. Despite previous findings of decreased interhemispheric structural connectivity in uCP, as measured with diffusion MRI in the corpus callosum (Pannek, Boyd, Fiori, Guzzetta, & Rose, 2014; Weinstein et al., 2014), functional connectivity does not seem to reflect these structural changes. Furthermore, research in functional connectivity in CP has typically assessed functional connectivity by means of a laterality index (Manning et al., 2015; Saunders et al., 2018), which does not resemble interhemispheric connectivity. Recently, Lee et al. combined both structural and functional measures in children with spastic diplegic CP to investigate the structure-function coupling, which was decreased in the patient population (D. Lee et al., 2017). They found that the efficiency of the functional motor network was decreased, despite a similar structural motor efficiency to controls (D. Lee et al., 2017). In this line, there is evidence of interhemispheric facilitation in uCP due to perinatal stroke instead of the typical interhemispheric inhibition, as measured with TMS (Eng, Zewdie, Ciechanski, Damji, & Kirton, 2018). However, we did not find differences in interhemispheric functional connectivity in any direction (from non-dominant M1 to contralateral ROIs, or vice versa) between TD and uCP, or between TD and CST wiring groups. As rsfMRI does not allow us to investigate facilitatory or inhibitory processes, we cannot reject that these processes may be different in each CST wiring group. Other advanced fMRI measures, like effective connectivity, may be needed to identify interhemispheric imbalance in the uCP population. Further research in uCP is needed to deduce causality, where we can infer the excitatory-inhibitory balance of individuals with uCP, measured for example with TMS, to better understand the specific pathophysiology of each CST wiring group.

Our **third hypothesis** investigated to what extent the sensorimotor network in the uCP group is more widespread than in controls, and the differences according to the CST wiring pattern, which was confirmed by the seed-to-voxel analysis. This analysis depicted higher connectivity in the total uCP group between the non-dominant M1 and the ipsilateral temporal lobe, and lower connectivity between the non-dominant M1 and both occipital poles, compared to controls. The temporal lobe is known to be important for semantic memory, and for this function, it is important that several brain regions participate in the comprehension of tasks (Binder & Desai, 2011). On the other hand, the occipital lobe is well known to be responsible for vision. The decreased connectivity seen between M1 and both occipital lobes in the uCP group may reflect an impaired visuomotor integration (Strigaro et al., 2015), as the communication between M1 and the visual network is very important for the motor and visual components of task performance (Eisenberg, Shmuelof, Vaadia, & Zohary, 2011). Secondly, the seed-to-voxel approach from the dominant M1 also identified a cluster covering sensory association areas where the functional connectivity was increased in the bilateral CST group compared to the other CST groups and the TD group. This may indicate a lack of functional specificity of the brain regions in the bilateral CST group, reflected in a larger and more extended network (Kanwisher, 2010). Areas that process distinct motor functions, as typically seen in the healthy brain, may be undistinguishable in this group due to the expanded sensorimotor network, as previously suggested by Burton et al. (Burton et al., 2009). In this line, an extended network may not be directly linked to a higher efficiency within the network (D. Lee et al., 2017), which may be the case in the bilateral CST wiring group.

The **fourth** and last **hypothesis** of this study was related to the combined impact that functional connectivity measures and the CST wiring pattern have on UL motor deficits in the uCP cohort. Although the different areas of the sensorimotor network included in this study are involved in motor execution and preparation, the connectivity of such a network at rest was barely related to deficits in grip strength, hand dexterity and bimanual performance in the whole uCP cohort. Also in the regression analysis, the identified differences in connectivity between groups did not significantly contribute to predict UL motor function, although there was an interesting trend indicating that higher connectivity in somatosensory association areas was related to poorer hand dexterity in combination with a bilateral or ipsilateral CST wiring, highlighting the importance of association and integration areas for UL function. However, the main predictor of UL motor deficits remains the underlying CST wiring, as we have shown in a recent study (Simon-Martinez, Jaspers, et al., 2018), and is also in agreement with previous literature (Staudt et al., 2004). The lack of a clear relation between functional connectivity from M1 and UL motor deficits shown in our study are in agreement with the recent findings of Saunders et al. (Saunders et al., 2018). Despite the low correlations found in theirs and our study, the potential value of functional connectivity in the uCP group may not be fully lost. It may be that the clinical tests do not reflect the specificity of the functional connectivity measures, as UL function was evaluated with scales that show an overall picture of the UL deficits, despite the fact that we had a fair representation of UL deficits (MACS levels I to III). Furthermore, the small sample size that we had in each CST wiring group may not have been enough to depict the potential impact of the functional connectivity on UL function in each group. On the contrary, it is plausible that functional connectivity in the uCP cohort due to white matter lesions does not serve as a biomarker on its own for this CP subgroup, but in other CP subgroups. There are surely other factors intermediating the complex relationship between functional connectivity and motor deficits. In this study, we included the CST wiring as previous literature highlighted its power in predicting UL deficits. However, the combination with other measures of microstructural integrity may give more accurate information. For example, a recent study showed that the decoupling between the structural and functional connectome may add information to understand the underlying pathophysiology of UL sensorimotor deficits (D. Lee et al., 2017). There is a clear need for multimodal neuroimaging studies in the uCP population, including different lesion types, to advance toward a more comprehensive understanding of the problems, which will lead to a more accurate definition of the targeted treatment.

### Strengths, limitations and future directions

This is the first study including a large sample of TD individuals as reference to investigate deficits in a moderate group of uCP participants with a homogeneous lesion type. In general, our results show very small within group variability in the functional connectivity measures of the TD cohort, suggesting that the connectivity measures of the sensorimotor network are quite replicable in TD children. However, the uCP cohort showed very large variability, suggesting that the study of functional connectivity may not be very sensitive in this population to be used as a biomarker, as other factors may influence the strength of the connectivity. Furthermore, the novel combination of functional connectivity measures and the underlying CST wiring pattern contributed to deepen our understanding of the pathophysiology of UL function in uCP.

Among the limitations of our study are that our results are not representative for other lesion types in uCP, such as cortico-subcortical lesions or malformations. These lesion types should be included in further research to provide a bigger picture of the distinct lesion mechanisms in uCP. Secondly, in this study we investigated cortico-cortical functional connectivity within the sensorimotor network, but also more remotely with other cortical areas. In line with our findings of a more widespread sensorimotor network, it seems interesting to further explore whereas cortico-subcortical connectivity or the connectivity among the subcortical structures (thalamus and striatum), which may shed light onto drawing the bigger picture of the impact of functional connectivity in uCP. Thirdly, despite the large sample size of the uCP cohort, the number of participants in each CST wiring group is still limited. Finally, we only included valid clinical measures that are widely used in uCP research to evaluate motor deficits. It would be of interest to also include other measures of sensory function, or even more specific measures (i.e. deficits in sensorimotor integration, visuomotor adaptation, motor learning…). Although sensory deficits are minimal in individuals with periventricular white matter lesions (Mailleux et al., 2017), including a more quantitative measure of sensory deficits may be interesting in future research. Similarly, more specific measures of sensorimotor integration of multisensory deficits (i.e. visuomotor adaptation) could be included, which could potentially be related to the aberrant functional connectivity of the sensorimotor network.

Future directions of functional connectivity in uCP should also be addressed to its potential of partially explaining the behavioural improvements after an intensive unimanual training such as constraint induced movement therapy, as the sensorimotor network becomes more lateralized (similar to healthy controls) (Manning et al., 2016). It has previously been shown that functional connectivity measured with the laterality index by rsfMRI, is a significant predictor of treatment response after constraint induced movement therapy in children with different lesion types (periventricular white matter and cortico-subcortical injuries), whereby an imbalanced sensorimotor network (i.e. stronger connectivity within the dominant hemisphere) predicts improvement in motor abilities after the treatment (Manning et al., 2015; Rocca et al., 2013). This highlights the plasticity of the resting motor network after treatment. Therefore, the potential value of functional connectivity of the sensorimotor network should be furthered explored as a predictor of treatment outcome and to understand the plastic changes that an intervention may implicate.

### Conclusion

Based on current study results, functional connectivity of the sensorimotor network at rest can identify connectivity patterns that are CST-dependent rather than specific from all the uCP population, in particular in the dominant hemisphere. Furthermore, functional connectivity seems to have little potential to predict UL motor deficits, as the type of CST wiring remains the main predictor of motor outcome. With this identification of functional connectivity features (higher connectivity in the dominant hemisphere and distinct pattern of remote connectivity), we hope to contribute to pave the way toward a better understanding of the underlying pathophysiology of UL function. Also, by identifying where the specific pathophysiology occurs, non-invasive brain stimulation protocols may be developed targeting these deficits while considering the underlying type of CST wiring pattern. Lastly, deeper knowledge of these characteristics may be also useful to delineate training programs or predicting treatment response in uCP.

## 9. Appendices

### Supporting Information

Table S1. MNI coordinates of the ROIs included in the analysis.

Table S2. Descriptive demographic data of each cohort.

Table S3. Correlation coefficients (Pearson’s r (p-value)) between functional connectivity measures and UL motor function in the uCP cohort.

## Acknowledgments

This work is funded by the Fund Scientific Research Flanders (FWO-project, grant G087213N) and by the Special Research Fund, KU Leuven (OT/14/127, project grant 3M140230). JB and FA have received funding from the Research Foundation Flanders (FWO), who received funding from the European Union’s Horizon 2020 research and innovation programme (European Research council, grant no 647047) to collect data of the typically developing individuals.

We would like to express our deepest gratitude to the children and families who participated in this study. We also specially thank Jasmine Hoskens for her help during the clinical assessments.

## References

Benjamini, Y., & Hochberg, Y. (1995). Controlling the false discovery rate: a practical and powerful approach to multiple testing. Journal of the Royal Statistical Society, 57(1), 289–300. https://doi.org/10.2307/2346101

Binder, J. R., & Desai, R. H. (2011). The neurobiology of semantic memory. Trends in Cognitive Sciences, 15(11), 527–536. https://doi.org/10.1016/j.tics.2011.10.001

Burton, H., Dixit, S., Litkowski, P., & Wingert, J. R. (2009). Functional connectivity for somatosensory and motor cortex in spastic diplegia. Somatosensory & Motor Research, 26(4), 90–104. https://doi.org/10.3109/08990220903335742

Chai, X. J., Castañán, A. N., Öngür, D., & Whitfield-Gabrieli, S. (2012). Anticorrelations in resting state networks without global signal regression. NeuroImage, 59(2), 1420–1428. https://doi.org/10.1016/j.neuroimage.2011.08.048

Davare, M., Andres, M., Cosnard, G., Thonnard, J.-L., & Olivier, E. (2006). Dissociating the role of ventral and dorsal premotor cortex in precision grasping. The Journal of Neuroscience : The Official Journal of the Society for Neuroscience, 26(8), 2260–8. https://doi.org/10.1523/JNEUROSCI.3386-05.2006

Di Martino, A., Yan, C. G., Li, Q., Denio, E., Castellanos, F. X., Alaerts, K.,…Milham, M. P. (2014). The autism brain imaging data exchange: Towards a large-scale evaluation of the intrinsic brain architecture in autism. Molecular Psychiatry, 19(6), 659–667. https://doi.org/10.1038/mp.2013.78

Eisenberg, M., Shmuelof, L., Vaadia, E., & Zohary, E. (2011). The Representation of Visual and Motor Aspects of Reaching Movements in the Human Motor Cortex. Journal of Neuroscience, 31(34), 12377–12384. https://doi.org/10.1523/JNEUROSCI.0824-11.2011

Eliasson, A.-C., Krumlinde-Sundholm, L., Rösblad, B., Beckung, E., Arner, M., Öhrvall, A.-M.,…Rosenbaum, P. (2006). The Manual Ability Classification System (MACS) for children with cerebral palsy: scale development and evidence of validity and reliability. Developmental Medicine and Child Neurology, 48(07), 549. https://doi.org/10.1017/S0012162206001162

Eng, D., Zewdie, E., Ciechanski, P., Damji, O., & Kirton, A. (2018). Interhemispheric motor interactions in hemiparetic children with perinatal stroke: Clinical correlates and effects of neuromodulation therapy. Clinical Neurophysiology, 129(2), 397–405. https://doi.org/10.1016/j.clinph.2017.11.016

Feys, H., Eyssen, M., Jaspers, E., Klingels, K., Desloovere, K., Molenaers, G., & De Cock, P. (2010). Relation between neuroradiological findings and upper limb function in hemiplegic cerebral palsy. European Journal of Paediatric Neurology : EJPN : Official Journal of the European Paediatric Neurology Society, 14(2), 169–77. https://doi.org/10.1016/j.ejpn.2009.01.004

Friston, K. J., Ashburner, J., Kiebel, S., Nichols, T., & Penny, W. D. (2007). Statistical parametric mapping: the analysis of funtional brain images. (K. Friston, J. Ashburner, S. Kiebel, T. Nichols, & P. William, Eds.). Amsterdam: Elsevier/Academic Press.

Gaberova, K., Pacheva, I., & Ivanov, I. (2018). Task-related fMRI in hemiplegic cerebral palsy-A systematic review. Journal of Evaluation in Clinical Practice, 24(4), 839–850. https://doi.org/10.1111/jep.12929

Genovese, C. R., Lazar, N. A., & Nichols, T. (2002). Thresholding of statistical maps in functional neuroimaging using the false discovery rate. NeuroImage, 15(4), 870–878. https://doi.org/10.1006/nimg.2001.1037

Gordon, A. M., Charles, J., & Wolf, S. L. (2006). Efficacy of constraint-induced movement therapy on involved upper-extremity use in children with hemiplegic cerebral palsy is not age-dependent. Pediatrics, 117(3), e363–73. https://doi.org/10.1542/peds.2005-1009

Gupta, D., Barachant, A., Gordon, A. M., Ferre, C., Kuo, H.-C., Carmel, J. B., & Friel, K. M. (2017). Effect of sensory and motor connectivity on hand function in pediatric hemiplegia. Annals of Neurology, 82(5), 766–780. https://doi.org/10.1002/ana.25080

Hinkle, D. E., & Wiersma, W. (1998). Applied Statistics for the behavioral sciences. Houghton Mifflin.

Holmefur, M., Aarts, P., Hoare, B., & Krumlinde-Sundholm, L. (2009). Test-retest and alternate forms reliability of the assisting hand assessment. Journal of Rehabilitation Medicine, 41(11), 886–891. https://doi.org/10.2340/16501977-0448

Holmström, L., Lennartsson, F., Eliasson, A. C., Flodmark, O., Clark, C., Tedroff, K.,…Vollmer, B. (2011). Diffusion MRI in corticofugal fibers correlates with hand function in unilateral cerebral palsy. Neurology, 77(8), 775–783. https://doi.org/10.1212/WNL.0b013e31822b0040

Holmström, L., Vollmer, B., Tedroff, K., Islam, M., Persson, J. K. E., Kits, A.,…Eliasson, A. C. (2010). Hand function in relation to brain lesions and corticomotor-projection pattern in children with unilateral cerebral palsy. Developmental Medicine and Child Neurology, 52(2), 145–152. https://doi.org/10.1111/j.1469-8749.2009.03496.x

Jeannerod, M., Arbib, M. A., Rizzolatti, G., & Sakata, H. (1995). Grasping objets: The cortical mechanisms of visuomotor transformation. Trends in Neurosciences, 18(7), 314–320. https://doi.org/10.1016/0166-2236(95)93921-J

Kanwisher, N. (2010). Functional specificity in the human brain: a window into the functional architecture of the mind. Proceedings of the National Academy of Sciences of the United States of America, 107(25), 11163–70. https://doi.org/10.1073/pnas.1005062107

Klingels, K., Demeyere, I., Jaspers, E., De Cock, P., Molenaers, G., Boyd, R., & Feys, H. (2012). Upper limb impairments and their impact on activity measures in children with unilateral cerebral palsy. European Journal of Paediatric Neurology, 16(5), 475–484. https://doi.org/10.1016/j.ejpn.2011.12.008

Krumlinde-Sundholm, L., & Eliasson, A.-C. C. (2003). Development of the Assisting Hand Assessment: A Rasch-built Measure intended for Children with Unilateral Upper Limb Impairments. Scandinavian Journal of Occupational Therapy, 10(1), 16–26. https://doi.org/10.1080/11038120310004529

Krumlinde-Sundholm, L., Holmefur, M., Kottorp, A., & Eliasson, A.-C. C. (2007). The Assisting Hand Assessment: current evidence of validity, reliability, and responsiveness to change. Developmental Medicine and Child Neurology, 49(4), 259–264. https://doi.org/10.1111/j.1469-8749.2007.00259.x

Lee, D., Pae, C., Lee, J. D., Park, E. S., Cho, S.-R., Um, M.-H.,…Park, H.-J. (2017). Analysis of structure-function network decoupling in the brain systems of spastic diplegic cerebral palsy. Human Brain Mapping, 38(10), 5292–5306. https://doi.org/10.1002/hbm.23738

Lee, J. D., Park, H. J., Park, E. S., Oh, M. K., Park, B., Rha, D. W.,…Park, C. Il. (2011). Motor pathway injury in patients with periventricular leucomalacia and spastic diplegia. Brain, 134(4), 1199–1210. https://doi.org/10.1093/brain/awr021

Louwers, A., Beelen, A., Holmefur, M., & Krumlinde-Sundholm, L. (2016). Development of the Assisting Hand Assessment for adolescents (Ad-AHA) and validation of the AHA from 18 months to 18 years. Developmental Medicine & Child Neurology, 58(12), 1303–1309. https://doi.org/10.1111/dmcn.13168

Mailleux, L., Klingels, K., Fiori, S., Simon-Martinez, C., Demaerel, P., Locus, M.,…Feys, H. (2017). How does the interaction of presumed timing, location and extent of the underlying brain lesion relate to upper limb function in children with unilateral cerebral palsy? European Journal of Paediatric Neurology : EJPN : Official Journal of the European Paediatric Neurology Society, 21(5), 763–772. https://doi.org/10.1016/j.ejpn.2017.05.006

Manning, K. Y., Fehlings, D., Mesterman, R., Gorter, J. W., Switzer, L., Campbell, C., & Menon, R. S. (2015). Resting State and Diffusion Neuroimaging Predictors of Clinical Improvements Following Constraint-Induced Movement Therapy in Children With Hemiplegic Cerebral Palsy. Journal of Child Neurology, 30(11), 1507–1514. https://doi.org/10.1177/0883073815572686

Manning, K. Y., Menon, R. S., Gorter, J. W., Mesterman, R. O. N. I. T., Campbell, C., Switzer, L., & Fehlings, D. (2016). Neuroplastic Sensorimotor Resting State Network Reorganization in Children with Hemiplegic Cerebral Palsy Treated with Constraint-Induced Movement Therapy. Journal of Child Neurology, 31(2), 220–226. https://doi.org/10.1177/0883073815588995

Mayka, M. A., Corcos, D. M., Leurgans, S. E., & Vaillancourt, D. E. (2006). Three-dimensional locations and boundaries of motor and premotor cortices as defined by functional brain imaging: A meta-analysis. NeuroImage, 31(4), 1453–1474. https://doi.org/10.1016/j.neuroimage.2006.02.004

Pannek, K., Boyd, R. N., Fiori, S., Guzzetta, A., & Rose, S. E. (2014). Assessment of the structural brain network reveals altered connectivity in children with unilateral cerebral palsy due to periventricular white matter lesions. NeuroImage: Clinical, 5, 84–92. https://doi.org/10.1016/j.nicl.2014.05.018

Papadelis, C., Ahtam, B., Nazarova, M., Nimec, D., Snyder, B., Grant, P. E., & Okada, Y. (2014). Cortical Somatosensory Reorganization in Children with Spastic Cerebral Palsy: A Multimodal Neuroimaging Study. Frontiers in Human Neuroscience, 8, 725. https://doi.org/10.3389/fnhum.2014.00725

Power, J. D., Barnes, K. A., Snyder, A. Z., Schlaggar, B. L., & Petersen, S. E. (2012). Spurious but systematic correlations in functional connectivity MRI networks arise from subject motion. NeuroImage, 59(3), 2142–2154. https://doi.org/10.1016/j.neuroimage.2011.10.018

Pruim, R. H. R., Mennes, M., Buitelaar, J. K., & Beckmann, C. F. (2015). Evaluation of ICA-AROMA and alternative strategies for motion artifact removal in resting state fMRI. NeuroImage, 112, 278–287. https://doi.org/10.1016/j.neuroimage.2015.02.063

Rocca, M. A., Turconi, A. C., Strazzer, S., Absinta, M., Valsasina, P., Beretta, E.,…Filippi, M. (2013). MRI Predicts Efficacy of Constraint-Induced Movement Therapy in Children With Brain Injury. Neurotherapeutics, 10(3), 511–519. https://doi.org/10.1007/s13311-013-0189-2

Saunders, J., Carlson, H. L., Cortese, F., Goodyear, B. G., & Kirton, A. (2018). Imaging functional motor connectivity in hemiparetic children with perinatal stroke. Human Brain Mapping. https://doi.org/10.1002/hbm.24474

Seghier, M. L. (2008). Laterality index in functional MRI: methodological issues. Magnetic Resonance Imaging, 26(5), 594–601. https://doi.org/10.1016/j.mri.2007.10.010

Simon-Martinez, C., Jaspers, E., Mailleux, L., Ortibus, E., Klingels, K., Wenderoth, N., & Feys, H. (2018). Corticospinal Tract Wiring and Brain Lesion Characteristics in Unilateral Cerebral Palsy: Determinants of Upper Limb Motor and Sensory Function. Neural Plasticity, 2018, 1–13. https://doi.org/10.1155/2018/2671613

Simon-Martinez, C., Mailleux, L., Ortibus, E., Fehrenbach, A., Sgandurra, G., Cioni, G.,…Klingels, K. (2018). Combining constraint-induced movement therapy and action-observation training in children with unilateral cerebral palsy: a randomized controlled trial. BMC Pediatrics, 18(1), 250. https://doi.org/10.1186/s12887-018-1228-2

Staudt, M. (2010). Reorganization after pre- and perinatal brain lesions. Journal of Anatomy, 217(4), 469–74. https://doi.org/10.1111/j.1469-7580.2010.01262.x

Staudt, M., Gerloff, C., Grodd, W., Holthausen, H., Niemann, G., & Krägeloh-Mann, I. (2004). Reorganization in congenital hemiparesis acquired at different gestational ages. Annals of Neurology, 56(6), 854–863. https://doi.org/10.1002/ana.20297

Staudt, M., Grodd, W., Gerloff, C., Erb, M., Stitz, J., & Krägeloh-Mann, I. (2002). Two types of ipsilateral reorganization in congenital hemiparesis: a TMS and fMRI study. Brain : A Journal of Neurology, 125, 2222–2237. https://doi.org/10.1093/brain/awf227

Strigaro, G., Ruge, D., Chen, J.-C., Marshall, L., Desikan, M., Cantello, R., & Rothwell, J. C. (2015). Interaction between visual and motor cortex: a transcranial magnetic stimulation study. The Journal of Physiology, 593(10), 2365–77. https://doi.org/10.1113/JP270135

Taylor, N., Sand, P. L., & Jebsen, R. H. (1973). Evaluation of hand function in children. Archives of Physical Medicine and Rehabilitation, 54(3), 129–35.

Theys, C., Wouters, J., & Ghesquière, P. (2014). Diffusion tensor imaging and resting-state functional MRI-scanning in 5- and 6-year-old children: training protocol and motion assessment. PloS One, 9(4), e94019. https://doi.org/10.1371/journal.pone.0094019

Tsao, H., Pannek, K., Fiori, S., Boyd, R. N., & Rose, S. (2014). Reduced integrity of sensorimotor projections traversing the posterior limb of the internal capsule in children with congenital hemiparesis. Research in Developmental Disabilities, 35(2), 250–260. https://doi.org/10.1016/j.ridd.2013.11.001

Van Dijk, K. R. A., Sabuncu, M. R., & Buckner, R. L. (2012). The influence of head motion on intrinsic functional connectivity MRI. NeuroImage, 59(1), 431–438. https://doi.org/10.1016/j.neuroimage.2011.07.044

Vandermeeren, Y., Davare, M., Duque, J., & Olivier, E. (2009). Reorganization of cortical hand representation in congenital hemiplegia. European Journal of Neuroscience, 29(4), 845–854. https://doi.org/10.1111/j.1460-9568.2009.06619.x

Weinstein, M., Green, D., Geva, R., Schertz, M., Fattal-Valevski, A., Artzi, M.,…Bashat, D. Ben. (2014). Interhemispheric and intrahemispheric connectivity and manual skills in children with unilateral cerebral palsy. Brain Structure and Function, 219(3), 1025–1040. https://doi.org/10.1007/s00429-013-0551-5

Whitfield-Gabrieli, S., & Nieto-Castanon, A. (2012). Conn : A Functional Connectivity Toolbox for Correlated and Anticorrelated Brain Networks. Brain Connectivity, 2(3), 125–141. https://doi.org/10.1089/brain.2012.0073

Zewdie, E., Damji, O., Ciechanski, P., Seeger, T., & Kirton, A. (2017). Contralesional Corticomotor Neurophysiology in Hemiparetic Children with Perinatal Stroke: Developmental Plasticity and Clinical Function. Neurorehabilitation and Neural Repair, 31(3), 261–271. https://doi.org/10.1177/1545968316680485

