## Supporting information (Tables S1-S3) for "Sensorimotor functional connectivity in unilateral cerebral palsy: influence of corticospinal tract wiring pattern and clinical correlates"

Table S1. MNI coordinates of the ROIs included in the analysis.

|  | **Non-dominant hemisphere** | | | **Dominant hemisphere** | | |
| --- | --- | --- | --- | --- | --- | --- |
|  | **x** | **y** | **z** | **x** | **y** | **z** |
| **PMd** | -30 | -7 | 63 | 30 | -7 | 63 |
| **PMv** | -51 | 4 | 24 | 51 | 4 | 24 |
| **M1** | -37 | -25 | 62 | 37 | -25 | 62 |
| **S1** | -40 | -27 | 53 | 40 | -27 | 53 |
|  | **Midline structure (x, y, z)** | | | | | |
| **SMA proper** |  | 0 | -10 | | 59 |  |

SMA, supplementary motor area; PMd, premotor cortex dorsal part; PMv, premotor cortex ventral part; M1, primary motor cortex; S1, primary sensory cortex.

Table S2. Descriptive demographic data of each cohort.

|  |  | **TD cohort (n=60)** | **Contralateral CST (n=9)** | **Bilateral CST (n=6)** | **Ipsilateral CST (n=9)** |
| --- | --- | --- | --- | --- | --- |
| **MACS levels** | n (%) |  |  |  |  |
| I |  |  | 7 (78) | 0 (0) | 1 (11) |
| II |  |  | 1 (11) | 5 (83) | 5 (56) |
| III |  |  | 1 (11) | 1 (17) | 3 (33) |
| **Age (years)** | mean (SD) | 14.54 (4.80) | 14.54 (4.18) | 10.88 (3.41) | 13.36 (5.11) |
| **Head motion (mean FD)**^†^ | mean (SD) | 0.26 (0.12) | 0.31 (0.14) | 0.47 (0.30) | 0.25 (0.14) |
| **Sex** | n (%) |  |  |  |  |
| Male |  | 46 (77) | 2 (22) | 3 (50) | 4 (44) |
| Female |  | 14 (23) | 7 (78) | 3 (50) | 5 (56) |
| **Dominant hand** | n (%) |  |  |  |  |
| Right |  | 54 (90) | 2 (22) | 5 (83) | 6 (67) |
| Left |  | 6 (10) | 7 (78) | 1 (17) | 3 (33) |

TD, typically developing; CST, corticospinal tract; MACS, Manual Ability Classification System; FD, frame wise displacement; SD, standard deviation. ^†^Head motion (mean FD) was not different between the TD and the uCP cohorts (p>0.05).

Table S3. Correlation coefficients (Pearson’s r (p-value)) between functional connectivity measures and UL motor function in the uCP cohort.

|  |  | **Bimanual performance (AHA)** | **Hand dexterity (JTHF test, log10)** | **Grip strength (log10)** |
| --- | --- | --- | --- | --- |
| **Intra FC Non-dom** | **M1-PMd** | 0.01 (0.97) | 0.06 (0.77) | -0.14 (0.50) |
|  | **M1-PMv** | 0.00 (0.99) | 0.07 (0.74) | -0.09 (0.69) |
|  | **M1-S1** | 0.20 (0.33) | -0.17 (0.41) | -0.14 (0.49) |
| **Intra FC Dom** | **M1-PMd** | 0.17 (0.42) | -0.06 (0.77) | -0.17 (0.42) |
|  | **M1-PMv** | -0.07 (0.75) | 0.17 (0.42) | 0.03 (0.88) |
|  | **M1-S1** | -0.21 (0.32) | 0.10 (0.65) | 0.00 (0.99) |
| **Intra FC Dom** | **M1-(PO-SMG)** | -0.06 (0.77) | 0.23 (0.18) | 0.07 (0.72) |
| **Inter FC Non-dom 🡪 Dom** | **M1-PMd** | -0.06 (0.78) | -0.02 (0.91) | 0.03 (0.89) |
|  | **M1-PMv** | 0.03 (0.89) | -0.07 (0.73) | -0.06 (0.78) |
|  | **M1-S1** | 0.01 (0.98) | -0.17 (0.42) | -0.06 (0.78) |
|  | **M1-SMA** | -0.30 (0.15) | 0.28 (0.18) | 0.02 (0.92) |
| **Inter FC Dom 🡪 Non-dom** | **M1-PMd** | 0.29 (0.15) | -0.20 (0.33) | -0.24 (0.25) |
|  | **M1-PMv** | -0.13 (0.54) | 0.09 (0.67) | 0.01 (0.95) |
|  | **M1-S1** | 0.35 (0.09) | -0.39 (0.05) | -0.36 (0.08) |
|  | **M1-SMA** | -0.12 (0.57) | 0.30 (0.15) | -0.21 (0.32) |
| **Inter** | **M1-M1** | -0.06 (0.77) | 0.28 (0.18) | 0.07 (0.72) |

FC, functional connectivity; Non-dom, non-dominant hemisphere; Dom, dominant hemisphere; M1, primary motor cortex; PMd, dorsal stream of the premotor cortex; PMv, ventral stream of the premotor cortex; S1, primary sensory cortex; SMA, supplementary motor area; PO, parietal operculum; SPM, supramarginal gyrus; AHA, assisting hand assessment; JTHF, Jebsen-Taylor hand function.
